# A specific agonist of the orphan nuclear receptor NR2F1 suppresses metastasis through the induction of cancer cell dormancy

**DOI:** 10.1101/2021.01.30.428967

**Authors:** Bassem D. Khalil, Roberto Sanchez, Tasrina Rahman, Carolina Rodriguez-Tirado, Stefan Moritsch, Alba Rodriguez Martinez, Brett Miles, Eduardo Farias, Mihaly Mezei, Julie F. Cheung, Ana Rita Nobre, Nupura Kale, Karl Christoph Sproll, Maria Soledad Sosa, Julio A. Aguirre-Ghiso

**Affiliations:** Division of Hematology and Oncology, Department of Medicine, New York, NY, USA; Department of Pharmacological Sciences, New York, NY, USA; Drug Discovery Institute, New York, NY, USA; Department of Otolaryngology, New York, NY, USA; Department of Oncological Sciences, New York, NY, USA; Black Family Stem Cell Institute, New York, NY, USA; Tisch Cancer Institute, Icahn School of Medicine at Mount Sinai, New York, NY, USA; Department of Oral, Maxillofacial and Plastic Facial Surgery, Medical Faculty, University Hospital of the Heinrich-Heine-University Düsseldorf, Düsseldorf, Germany

## Abstract

Disseminated tumor cells (DTCs) in secondary organs often remain dormant for a long period of time before re-awakening and growing into overt metastases. We have previously identified NR2F1/COUP-TF1, an orphan nuclear receptor, as a master regulator of tumor cell dormancy in head and neck squamous cell carcinoma (HNSCC) and other cancer types. Here we describe the identification and function of a novel NR2F1 agonist herein referred to as compound 26 (C26). C26 was found to specifically activate NR2F1 in HNSCC cells, leading to increased NR2F1 transcription and nuclear protein accumulation. C26-mediated activation of NR2F1 induced growth arrest of HNSCC PDX line and cell lines in 3D cultures *in vitro* and on chicken embryo chorioallantoic membrane (CAM) *in vivo*. The effect of C26 on growth arrest was lost when NR2F1 was knocked out by CRISPR/Cas9. C26-induced growth arrest was mediated by activation of an NR2F1-regulated dormancy program, including upregulation of cyclin-dependent kinase (CDK) inhibitor p27 and the transcription factors retinoic acid receptor β (RARβ) and Sox9. In mice bearing HNSCC PDX tumors, combined adjuvant and neo-adjuvant treatment with C26 resulted in complete inhibition of lethal lung metastasis. Mechanistic analysis showed that lung DTCs in C26-treated mice displayed an NR2F1^hi^/p27^hi^/Ki-67^lo^ phenotype, which kept them dormant in a single-cell state preventing their outgrowth into overt metastases. Our work reveals a novel NR2F1 agonist and provides a proof of principle strategy supporting that inducing DTC dormancy using NR2F1 agonists could be used as a therapeutic strategy to prevent metastasis.

## Introduction

Metastasis is the leading cause of cancer related deaths and arises from disseminated tumor cells (DTCs) that seed secondary organs. These DTCs can remain in a dormant state for years or decades before growing into symptomatic overt metastases [1]. This period of latency occurs in multiple types of cancers and it is not limited to distant organs but also to loco-regional recurrences, a problem that is particularly significant in head and neck squamous cell carcinoma (HNSCC). More than half the patients with advanced disease develop recurrences that are usually not responsive to conventional therapies [2]. This highlights the need for interventional therapies that prevent reactivation of dormant DTCs after treatment of primary tumors, a window of treatment opportunity that is currently missed.

Our laboratory has focused on understanding the mechanisms that govern tumor cell dormancy. We identified the nuclear receptor subfamily 2 group F member 1 (NR2F1; also known as COUP-TF1) as a master regulator of this process [3]. NR2F1 is a nuclear receptor of the steroid/thyroid hormone receptors superfamily and is an orphan receptor. It has a DNA binding domain (DBD) consisting of two conserved Zinc-finger motifs and a ligand-binding domain (LBD) [4]. It regulates transcription either directly by binding as a dimer to direct repeats (DR) on DNA and recruiting co-activator or co-repressor complexes or indirectly by acting as a co-factor to other nuclear receptors [5]. Depending on the context, NR2F1 is able to activate or repress transcription of effector genes [5]. NR2F1 also plays an epigenetic role in mediating global histone modifications by interacting with or recruiting chromatin-remodeling enzymes [5].

We found that NR2F1 is epigenetically silenced in proliferating cancer cells and it is upregulated in dormant residual HNSCC cells in a PDX model and in prostate cancer DTCs isolated from patients [3]. In HNSCC patients, NR2F1 expression is absent or low in HNSCC primary tumors, recurrent tumors, and metastases compared to benign adjacent oral mucosa [3]. When upregulated, NR2F1 induces expression of a dormancy gene signature, including the transcription factors SOX9 and retinoic acid receptor β (RARβ). These factors activate expression of cyclin-dependent kinase (CDK) inhibitors p27 and p16, which leads to G0/G1 cell cycle arrest and cell quiescence [3]. Surprisingly, the NR2F1-associated signature was also found enriched in ER^+^ breast cancer tumors, and patients carrying primary lesions enriched for this signature showed delayed time to metastasis [6]. More specific to DTC biology, we also reported that breast cancer patients who carried in their bone marrow DTCs positive for NR2F1 were less likely to develop and die from bone metastasis compared to those negative or low for NR2F1 [7]. NR2F1 is also associated with alterations in a breast cancer susceptibility locus (Mcs1) [8]. These data argue that NR2F1 is a strong negative regulator of breast cancer, HNSCC, and other cancers, and that a unique function of NR2F1 is the induction of cancer cell dormancy, likely a role related to its lineage commitment function in development [9, 10].

ShRNA-mediated downregulation of NR2F1 results in reactivation of dormant HNSCC tumor cells leading to loco-regional relapse in surgery margins and development of lung and splenic metastasis in mice [3]. This shows that NR2F1 plays a critical role in initiating and maintaining tumor cell dormancy, a biological function that has been reproduced by independent labs in various cancer models [3, 6, 11–13]. We also showed that an epigenetic reprogramming therapy based on the combination of low dose 5-azacytidine and retinoic acid resulted in an upregulation of NR2F1 and dormancy induction in various cancer models [3]. While NR2F1 is silenced, it is not completely absent in many tumors [3, 6]. Additionally, although not directly tested, NR2F1 is predicted to regulate its own expression based on computational analysis and chromatin status data [3, 14]. Hence, experimental and clinical data support that activating NR2F1 using a small molecule agonist represents an attractive clinical strategy to induce dormancy and prevent recurrence and metastasis by restoring its expression and function in DTCs. Here we report the discovery of an NR2F1 selective agonist that can induce and activate NR2F1 leading to the powerful activation of a dormancy program in cancer cells. Importantly, use of this agonist in a preclinical neo-adjuvant and adjuvant setting completely suppressed macro-metastatic growth *via* the induction of DTC dormancy. Our work reveals a proof of principle strategy to limit metastatic growth by activation of dormancy mechanisms.

## Results

### Modeling of the NR2F1 ligand-binding domain identifies NR2F1 agonists in silico

There is no available crystal structure of NR2F1 or its ligand-binding domain (LBD). However, we were able to build a structural model based on homologous proteins of known structure. Among the proteins of known structure, the most similar to NR2F1 is NR2F2 [15]. However, the available NR2F2 structure is in the auto-repressed conformation, in which the agonist-binding site is occluded. Hence, a model of NR2F1 ligand binding domain (LBD) was built using as template the structure of retinoic acid receptor alpha (RXRα) in complex with 9-cis retinoic acid (PDB code 1fm6, ~40% sequence identify to NR2F1 over the LBD) [16]. The resulting model of NR2F1 LBD represents its active conformation and was used as the target for virtual screening (**Fig.1A**). This led to a structure-based virtual screening approach to identify small molecules with the potential to act as agonists of NR2F1.

**Figure 1.**
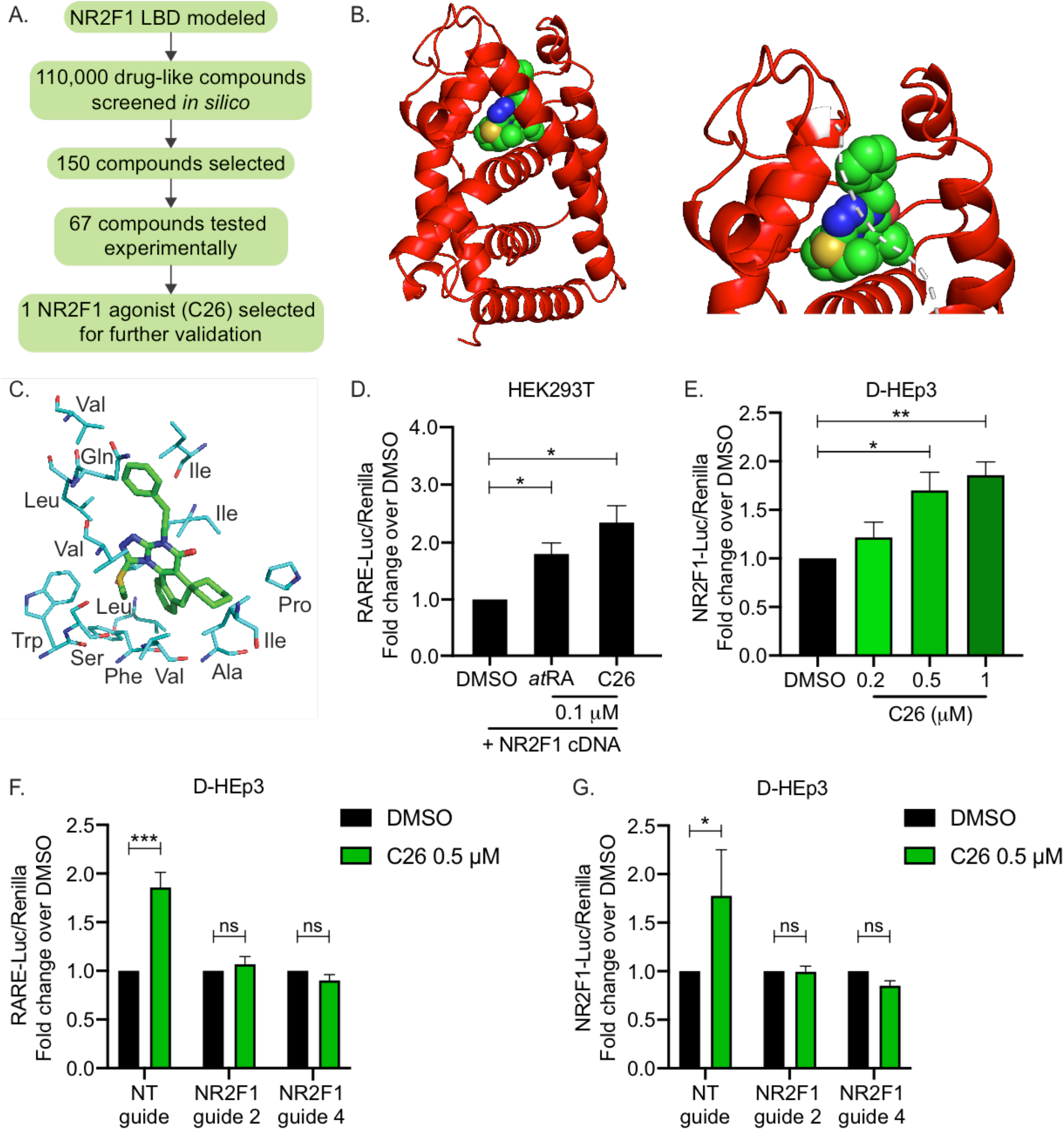
NR2F1 LBD modeling and agonist screen. **A.** Diagram depicting the approach used to screen for NR2F1 agonists. **B.** Left panel: ribbon diagram (in red) of NR2F1 LBD modeled using MODELLER v9.10 shown with the agonist C26 (sphere representation) docked in the binding site. Right panel: close-up of C26 docked in binding site; white dotted line represents part of the helix that was removed for better view of the binding site. **C.** Stick representation showing interaction of C26 with mostly hydrophobic residues in the biding site of NR2F1 LBD. **D.** Graph shows fold change of Luciferase/Renilla activity using RARE-luciferase reporter system in HEK293T cells with NR2F1 overexpression. Cells were treated with DMSO, *all trans* retinoic acid (*at*RA) (0.1 μM), or C26 (0.1 μM). Data represent mean +/− SEM from 3 independent experiments. * p<0.05 by *t test*. **E.** Graph shows fold change of Luciferase/Renilla activity using NR2F1-luciferase reporter system in D-HEp3 cells. Cells were treated with DMSO or C26 (0.2, 0.5, or 1 μM). Data represent mean +/− SEM from 3 independent experiments. * p<0.05, ** p<0.01 by ANOVA. **F and G.** Graphs show fold change of Luciferase/Renilla activity using RARE-luciferase (F) or NR2F1-luciferase (G) reporter systems in control D-HEp3 expressing non-targeting gRNA (NT gRNA) or two D-HEp3 cell lines with NR2F1 konckout using two separate guide RNAs (guide 2 and guide 4). Cells were treated with DMSO or C26 (0.5 μM). Data represent mean +/− SEM from 3 independent experiments. * p<0.05; *** p<0.001; ns: not significant by ANOVA.

Autodock and eHITS programs were used to screen a library of 110,000 drug-like compounds. The top 50 compounds identified by each program (based on docking score) were combined with the top 50 compounds ranked by the sum of their Autodock4 and eHits ranks. All selected compounds were visually inspected in the context of the target-binding site to select the most promising ones. Of the 150 initial compounds, 67 were selected for experimental validation using a retinoic acid response element (RARE) luciferase reporter system in HEK293T cells with NR2F1 overexpression (data not shown).

Of the 67 compounds, one agonist herein referred to as compound C26 (C26), consistently and significantly resulted in RARE reporter activation and was thus chosen for further validation. A model of NR2F1 in complex with C26, generated using the methods described above (**Fig.1B**), shows that C26 interacts with mostly hydrophobic residues in the biding pocket in NR2F1 (**Fig.1C**). C26 was found to induce luciferase expression from RARE reporter by 2.4-fold, a level that is comparable to 100 nM *all trans* retinoic acid (*at*RA) (**Fig.1D**), which activates transcription of target genes with RARE *via* binding to retinoic acid receptors (RARs) [17]. Since the RARE-luciferase bioassay reports on the combined signaling of RARs and NR2F1 as a co-activator [18], we used another reporter system where luciferase expression is driven by NR2F1 *cis-*regulatory element in dormant HEp3 (D-HEp3) HNSCC cells that express high endogenous levels of NR2F1 [3, 14]. Results show that C26 treatment significantly induces luciferase expression by 1.7 and 1.9-fold at 0.5 and 1 μM, respectively (**Fig.1E**), further confirming that C26 acts as an NR2F1 agonist. To verify that the effect of C26 on NR2F1-luciferase reporter activation is specific, we generated D-HEp3 cell lines where NR2F1 is knocked out by CRISPR/Cas9 using two different guide RNAs (gRNAs), which was confirmed by western blot (**Fig.S1A**). C26 effect on the RARE- and NR2F1-luciferase reporters in the NR2F1 knockout cells along with a D-HEp3 cell line expressing a non-targeting (NT) gRNA was evaluated. While C26 activates both reporters in control cells expressing NT gRNA, the effect of C26 is completely abrogated in NR2F1 knockout cell lines (**Fig.1F and G**). These results indicate that the effect of C26 in cells is dependent on the presence of NR2F1 and confirm the on-target effect of C26. We next assessed whether C26 has the ability to bind and activate RXRα, whose structure was used to model NR2F1 LBD, using time-resolved fluorescence resonance energy transfer (TR-FRET) RXRα co-activator assay *in vitro*. Importantly, C26 does not activate RXRα even at concentrations that are 5 to 10-fold above those that we use in our assays (**Fig.S1B**). These results further support that C26 is a selective agonist for NR2F1 in human cancer cells.

### C26 upregulates NR2F1 and a downstream dormancy program

In a previous study, we showed that knockdown of NR2F1 reduces repressive chromatin marks on its own promoter allowing an active chromatin state [3]. Additionally, computational identification of a transcription factor network active in dormant HNSCC cells revealed that NR2F1 is a central node, and it was predicted to regulate its own expression [14]. These observations suggest that upon its activation, NR2F1 might upregulate its own expression, and that even NR2F1^lo^ tumor cells could be activated to re-express this nuclear receptor. This hypothesis has not been tested before due to the lack of experimental tools to activate NR2F1. To test this possibility, we used tumorigenic HEp3 (T-HEp3) cells, a highly proliferative HNSCC PDX line that was obtained from a lymph node metastasis with primary carcinoma in the buccal mucosa [19] and maintained *in vivo*. These cells express low but detectable levels of NR2F1 [3, 12]. T-HEp3 cells were pre-treated with DMSO or 0.5 μM C26 and inoculated *in vivo* on chicken embryo chorioallantoic membrane (CAM) with continuous daily treatments. After 7 days, tumors were excised and mRNA was extracted. qPCR analysis revealed that C26 treatment results in 2.3-fold increase in NR2F1 mRNA levels in tumors (**Fig.2A**). To evaluate if this increase in mRNA also leads to upregulation of NR2F1 at the protein level, we performed a similar *in vivo* CAM experiment, but this time the tumors were dissociated and cytospins of cells were immunostained for NR2F1. T-HEp3 cells are distinguished from avian cells by immunostaining for vimentin, an intermediate filament that is abundantly expressed by HEp3 cells and has been previously used to detect these cells [3, 12, 20]. Indeed, the percentage of NR2F1^+^ cells is dramatically increased from around 4% in DMSO control tumors, which is consistent with what was previously reported [3] to around 42% in tumors treated with C26 (**Fig.2B and 2C**). This was determined using a stringent mask for strong nuclear NR2F1 signal, which appears as prominent clusters that we had reported previously in HNSCC and breast cancer DTCs [3, 7, 12]. These data support that the orphan receptor is upregulated, and that it is located in the nucleus upon C26 treatment. To further confirm the effect of C26 on NR2F1 expression, we used an *in vitro* 3D assay, where T-HEp3 cells were plated at low density in Matrigel. Cells were treated with DMSO or C26 (at 0.2, 0.5, and 1 μM) for 4 days. Immunostaining of fixed cells in the 3D Matrigel assay showed that NR2F1 protein level is indeed upregulated with C26 treatment. This is evident by a significant increase in the mean fluorescent intensity (MFI) of nuclear NR2F1 in C26-treated cells (at 0.2, 0.5, and 1 μM) compared to control cells treated with DMSO (**Fig.2D and 2E**). These data confirm that NR2F1 activation by C26 treatment results in upregulation of NR2F1 mRNA and nuclear protein levels.

**Figure 2.**
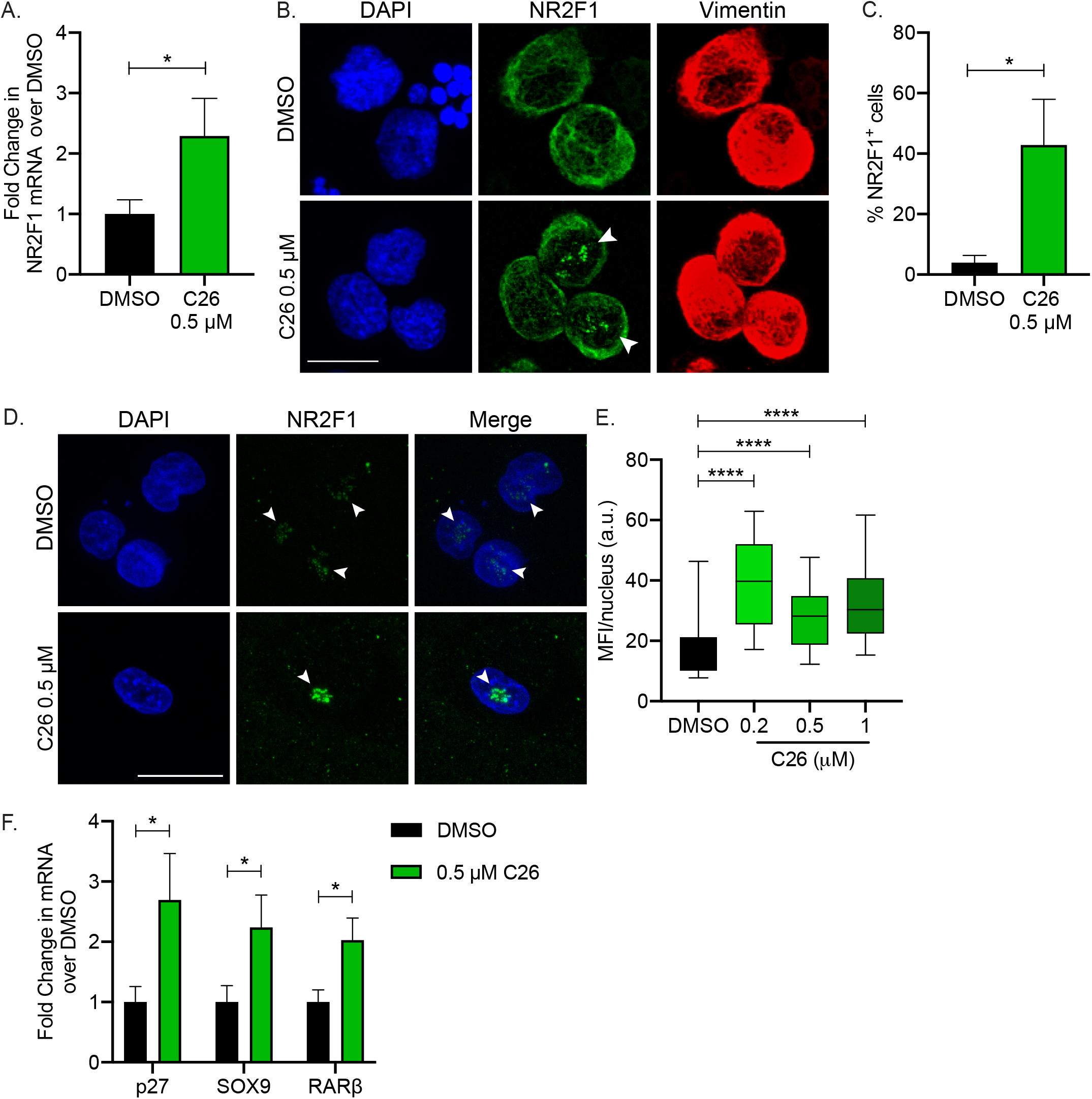
C26 upregulates NR2F1 expression. **A.** T-HEp3 cells pre-treated with DMSO or C26 (0.5 μM) were inoculated on CAM. After 7 days, tumors were collected and RNA extracted. Graph shows fold difference in NR2F1 mRNA levels obtained by qPCR and normalized to DMSO. Data represent mean +/− SEM from 4 tumors per group. * p<0.05 by *t test*. **B and C.** DMSO or C26 (0.5 μM) treated CAM tumors were dissociated and cell cytospins were generated. Cytospins were immunostained for NR2F1 and nuclei counterstained with DAPI. The percentage of cells with positive nuclear NR2F1 signal was quantitated. Scale bar: 10 μm; arrowheads indicate nuclear NR2F1. Graph shows mean % NR2F1^+^ cells +/− SEM from 4 tumors per group (87 cells for DMSO and 128 cells for C26). * p<0.05 by *t test*. **D and E.** T-HEp3 cells were plated in Matrigel and treated with DMSO or C26 (0.5 μM). After 4 days, cells were fixed and immunostained for NR2F1. Scale bar: 50 μm; arrowheads indicate nuclear NR2F1. Graph shows box (25 to 75 percentile) and whiskers (minimum to maximum values) of nuclear NR2F1 mean fluorescence intensity (MFI) per cell (64 cells for DMSO, 26 cells for 0.2 μM, 29 cells for 0.5 μM, 23 cells for 1 μM from two independent experiments). **** p<0.0001. **F.** SOX9, RARβ, and p27 mRNA levels were measured using qPCR in DMSO or C26 treated CAM tumors described in A. Graph shows fold difference in mRNA levels obtained by qPCR and normalized to DMSO. Data represent mean +/− SEM from 4 tumors per group. * p<0.05 by *t test*.

We had shown that NR2F1 induces tumor cell quiescence by binding to the promoters and inducing the expression of the transcription factors SOX9 and RARβ and also by upregulating the cyclin-dependent kinase (CDK) inhibitor p27 [3]. To determine if activation of NR2F1 by C26 and its nuclear accumulation was accompanied by an increase in NR2F1 target gene expression, we measured the effect of C26 treatment on mRNA levels of these downstream components in *in vivo* CAM tumors. Indeed, mRNA levels of SOX9, RARβ, and p27 are all significantly upregulated in C26-treated tumors compared to DMSO control (**Fig.2F**). Interestingly, C26 treatment has no effect on mRNA level of DEC2 (**Fig.S2**), a transcription factor that in dormant HNSCC and breast cancer cells is primarily regulated by transforming growth factor-β2 (TGF-β2) signaling [20]. Hence, C26 does not affect the TGF-β2-DEC2 pathway, which has been shown to mediate dormancy signals in a manner that is parallel to and independent from NR2F1 [3]. Collectively, these results show that C26 treatment induces NR2F1 expression and nuclear accumulation, and it also selectively activates canonical NR2F1-driven dormancy pathway genes.

### C26 induces growth arrest via NR2F1

We have previously shown that NR2F1 controls a cancer cell quiescence program in HNSCC and other cancer cells [3]. To determine if the increase in NR2F1 and the activation of NR2F1-dependant dormancy pathway described above leads to growth arrest, we plated single T-HEp3 cells at low density in Matrigel to mimic solitary DTC biology and treated daily with DMSO or C26 (at 0.2, 0.5, or 1 μM). After 4 days, number of single cells or colonies (defined as cell clusters with 3 or more cells) was determined by counting under light microscope. Results show that C26 treatment keeps cells in a single cell stage as evident by the higher percentage of single cells (**Fig.3A**) and lower percentage of clusters (**Fig.3B**) compared to DMSO. Accordingly, staining for the proliferation marker Ki-67 in the Matrigel assay shows that C26 significantly decreases the percentage of Ki-67^+^ proliferating T-HEp3 cells (**Fig.3C and S3A**). Similar results were obtained in two separate HNSCC cell lines, FaDu and SQ20B, where C26 treatment resulted in significantly higher percentage of single cells (**Fig.3D and 3E**) and lower percentage of Ki-67^+^ proliferating cells (**Fig.3F and 3G)**compared to DMSO. We then evaluated the effect of C26 on cell growth *in vivo* in the CAM model. T-HEp3 cells were pre-treated with DMSO or 0.5 μM C26 for 4-5 days in culture then inoculated on CAM, with or without continuous treatment. After 7 days, tumors were excised and dissociated using collagenase treatment, and the cell numbers were counted. Results show that C26 inhibits tumor growth on CAM both with pre-treatment only (**Fig.3H**) and to a higher extent with continuous treatment (**Fig.3I**). Importantly, staining for cleaved caspase-3 revealed that >99% of cells in both DMSO and C26 treated groups are negative for this apoptosis marker indicating that C26 treatment is not inducing cell death (**Fig.S3B**). The fact that pre-treatment with C26 alone was able to suppress growth a week later suggests that NR2F1 activation may be inducing a reprogramming that self-sustains the dormant phenotype.

**Figure 3.**
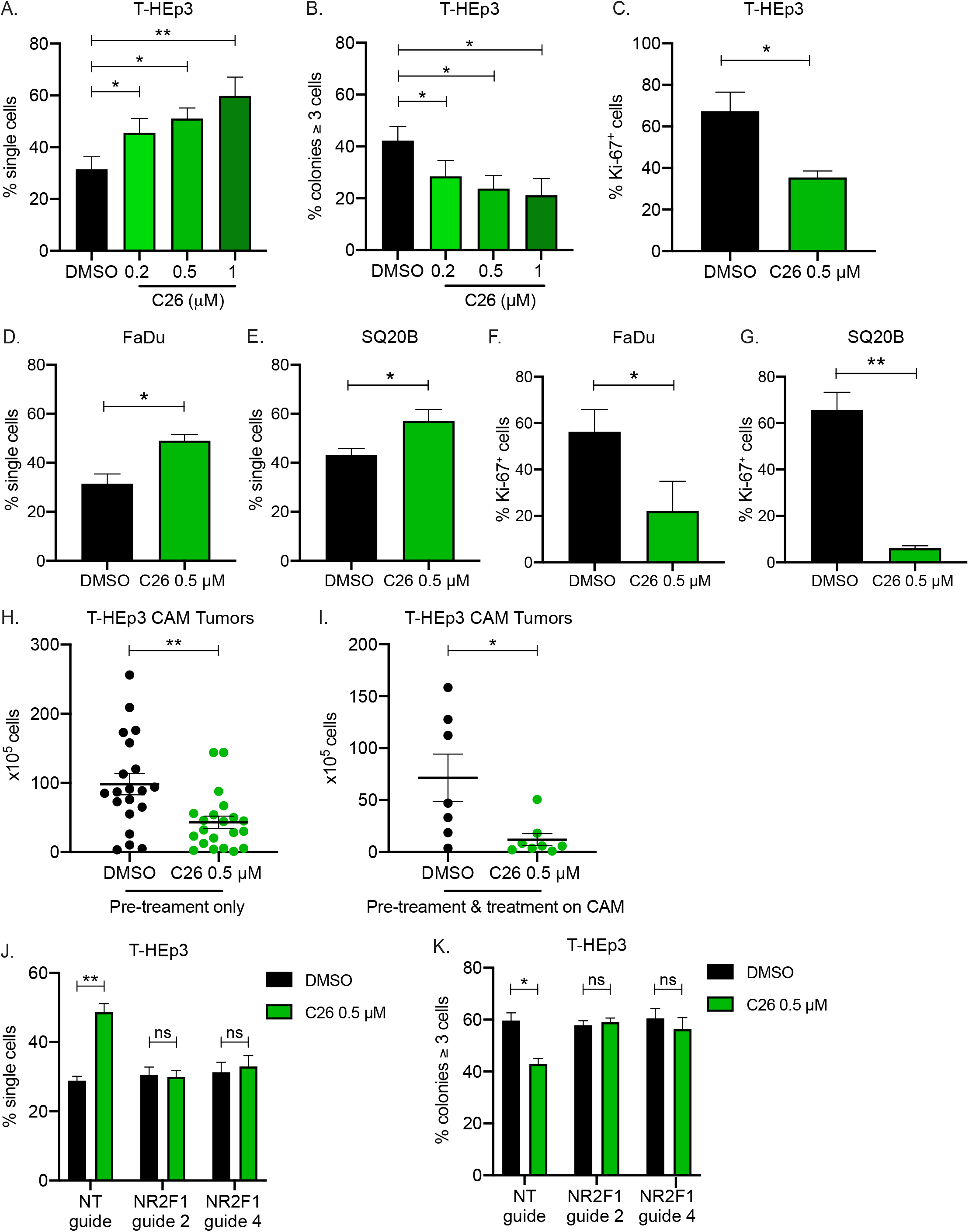
C26 induces growth arrest via NR2F1. **A and B.** T-HEp3 cells were plated in Matrigel and treated with DMSO or C26 (0.2, 0.5, or 1 μM). After 4 days, cells were counted manually under a microscope. Graphs show percentage of single cells (A) or colonies with 3 or more cells (B). Data represent mean +/− SEM from 3 independent experiments. * p<0.05; **P<0.01 by ANOVA. **C.** T-HEp3 cells were plated in Matrigel and treated with DMSO or C26 (0.5 μM). After 4 days, cells were fixed and immunostained for Ki-67. Graphs show percentage of Ki-67^+^ cells. Data represent mean +/− SEM from 3 independent experiments. * p<0.05 by *t test*. **D and E.** FaDu (D) or SQ20B (E) cells were plated in Matrigel and treated with DMSO or C26 (0.5 μM). After 4 days, cells were counted manually under a microscope. Graph shows percentage of single cells. Data represent mean +/− SEM from 3 independent experiments. * p<0.05 by *t test*. **F and G.** FaDu (F) or SQ20B (G) cells were plated in Matrigel and treated with DMSO or C26 (0.5 μM). After 4 days, cells were fixed and immunostained for Ki-67. Graphs show percentage of Ki-67^+^ cells. Data represent mean +/− SEM from 3 independent experiments. * p<0.05; ** p<0.01 by *t test*. **H.** T-HEp3 were pre-treated in culture for 5-7 days with DMSO or C26 (0.5 μM) and inoculated on CAM without continuous treatment. After 7 days, tumors were collected and dissociated and cell number per tumor was determined. Graph shows mean number of cells per tumor +/− SEM from 20 tumors (DMSO) and 21 tumors (C26). ** p<0.01 by *t test*. **I.** T-HEp3 were pre-treated in culture for 5-7 days with DMSO or C26 (0.5 μM) and inoculated on CAM with continued daily treatment. After 7 days, tumors were collected and dissociated and cell number per tumor was determined. Graph shows mean number of cells per tumor +/− SEM from 7 tumors (DMSO) and 8 tumors (C26). * p<0.05 by *t test*. **J and K.** T-HEp3 cells with non-targeting (NT) gRNA or cells where NR2F1 was knocked out by CRISPR using two different NR2F1 gRNAs (NR2F1 g2 and g4) were plated in Matrigel and treated with DMSO or C26 (0.2, 0.5, or 1 μM). After 4 days, cells were counted manually under a microscope. Graphs show mean number of single cells (J) or colonies with more 3 or more cells (K). Data represent mean +/− SEM from 2 independent experiments. * p<0.05; ** p<0.01 by ANOVA.

To confirm that the effect of C26 on cell growth is mediated by NR2F1, we generated T-HEp3 cell lines where NR2F1 is knocked out by CRISPR/Cas9 using two different guide RNAs (gRNAs) and confirmed the knockout by western blot (**Fig.S3C**). C26 effect on these cells lines, along with a T-HEp3 cell line expressing a non-targeting (NT) gRNA, was evaluated in the previously described 3D Matrigel assay. While C26 increases the percentage of single cells and decreases the number of clusters in cells with NT gRNA in a manner similar to the parental cell line, the effect of C26 was completely abrogated in the two NR2F1 knockout cell lines (**Fig.3J and 3K**). These results along with the *in vivo* detection of dormancy genes detected in Fig.2F and the selectivity controls showed in Fig.1 indicate that the growth suppressive effects of C26 in cancer cells is dependent on an NR2F1-driven dormancy program and confirms the on-target effect of C26.

### C26 inhibits primary tumor growth and metastatic growth in lungs

The above experiments establish that C26 activates NR2F1 and induces dormancy of HNSCC cells in *in vitro* 3D models and *in vivo* on the CAM. However, the CAM system does not allow to monitor long-term phenotypes in target organs since the tumors can be grown for a maximum of 7 days. Thus, we assessed whether C26-mediated activation of NR2F1 induces tumor cell dormancy and inhibits metastatic growth in mice. *In vivo* propagated GFP-tagged T-HEp3 PDX cells were injected subcutaneously in the interscapular region of Balb/c nu/nu mice. Tumors were allowed to develop and reach 300 mm^3^ in volume, after which neo-adjuvant treatment with either DMSO or C26 (0.5 mg/kg/day) was administered intra-peritoneally for 5 days. After two days of rest, tumors were surgically removed and four cycles of adjuvant treatment (5 days of treatment followed by two days of rest – 28 days total) were administered (**Fig.4A**). Excised tumors were weighed at the time of surgery and banked, and then lungs were collected when the mice were sacrificed at the end of the adjuvant treatment period. One lung lobe was processed for collagenase dissociation while other lobes were prepared for formalin fixation and paraffin embedding. Unexpectedly, C26 treatment inhibited primary tumor growth in 8 out 12 mice (**Fig.4B**). Importantly, analysis of the total number of GFP^+^ tumor cells per lung lobe after collagenase dissociation revealed that, without distinguishing between solitary DTCs or large or small metastatic lesions frequency, all mice that were treated with C26 had almost half the number of tumor cells compared to DMSO-treated mice (**Fig.4C**). This metastasis inhibitory effect of the NR2F1 agonist is independent of the effect on primary tumor growth since primary tumor weight and number of lung DTCs are not correlated (**Fig.S4**). Notably, there were no detectable negative side effects during C26 treatment or at autopsy.

**Figure 4.**
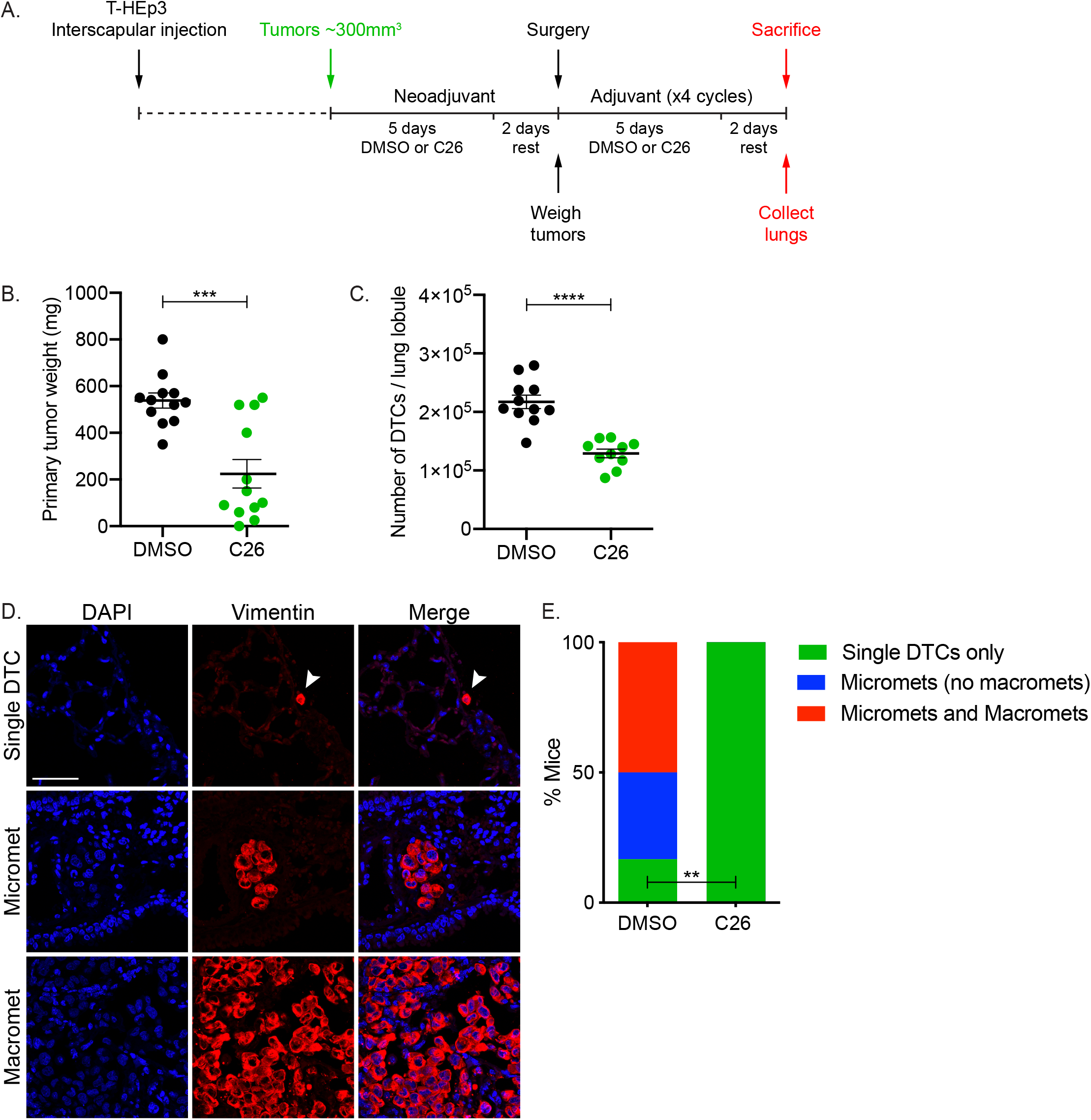
C26 suppresses primary tumor growth and lung metastatic growth in mice. **A.** Schematic depicting xenograft establishment as well as neoadjuvant and adjuvant treatment with DMSO or C26 (0.5 mg/kg/day) in mice. **B.** Graph shows weight of primary tumors surgically resected after the neoadjuvant phase. Data represent mean +/− SEM from 12 mice per group. *** p<0.001 by *t test*. **C.** Graph shows number of T-HEp3-GFP^+^ cells isolated from collagenase-digested lung lobules. Cells were counted under a fluorescence microscope and the total number of cells per lobule was calculated. Data represent mean +/− SEM from 11 mice (DMSO) and 10 mice (C26). **** p<0.0001 by *t test*. **D**. Whole lung lobules from experiment described in A. were immunostained for vimentin to detect single DTCs, micrometastases (<50 cells), or macrometastases (>50 cells). Scale bar: 50 μm. **E.** Graph shows percentage of DMSO or C26 treated mice with single DTCs, micrometastases only, or micrometastases and macrometastases. ** p<0.01 by Fisher’s exact test.

To provide more granularity on the metastases phenotypes, we immunostained lungs using an antibody for vimentin, an intermediate filament that is abundantly expressed in T-HEp3 cells and has been previously used to detect these cells in lungs [3, 12, 20]. We then determined the number of lungs with solitary cells only, micrometastases (<50 cells), and macrometastases (>50 cells) (**Fig.4D**). This analysis showed that 33% and 50% of DMSO-treated mice have micrometastases only or micrometastases and macrometastases in their lungs, respectively. However, 0% of mice treated with C26 showed micro or macro-metastases, and only single DTCs persisted in the lungs of these mice (**Fig.4E**). We have previously reported that in this aggressive model of HNSCC and in the absence of NR2F1, solitary DTCs in lungs transition out of a short-term quiescence and form metastasis [3]. Our current data indicates that while tumor cells were able to disseminate to the lungs and lodge in a solitary cell state as we showed previously [20], C26 treatment prevented single cells from dividing and growing into overt metastases. This is consistent with what we found *in vitro* in the 3D Matrigel cultures, where C26 treatment arrested cells in a single cell state.

### C26 treatment suppresses lung metastasis by activating an NR2F1^hi^/p27^hi^/Ki67^lo^ dormancy profile in solitary DTCs

We next sought to determine if the inability of lung DTCs to divide and grow into overt metastases in C26-treated mice is due to induction of NR2F1 and a dormancy phenotype. However, we first aimed to investigate the clinical relevance of NR2F1 by determining whether NR2F1 could pinpoint solitary DTCs, which are presumed to be dormant, in HNSCC patients [21]. To address this, we stained patient samples for NR2F1 in normal adjacent epithelium, primary tumors, and lymph nodes that were confirmed to have solitary HNSCC DTCs but not over metastasis [21]. Pancytokeratin was used as a marker to identify cells of epithelial origin in all tissues [22]. Cells were scored as NR2F1^hi^, NR2F1^med^, or NR2F1^−/low^. This analysis revealed that the percentage of NR2F1^hi^ cells drops from 28% in normal epithelium to 2.4% in primary tumors, while 20% of solitary DTCs are NR2F1^hi^. Conversely, the percentage of NR2F1^−/low^ is 39% in normal epithelium, 84% in primary tumors, and 63% in solitary DTCs (**Fig.5A and 5B**). These results indicate that solitary DTCs are more frequently positive for NR2F1 than cells in primary tumors, supporting that NR2F1 could mark dormant solitary DTCs in HNSCC patients. We had already confirmed in a previous report that NR2F1 expression is silenced in overt HNSCC lymph node metastasis [3].

**Figure 5.**
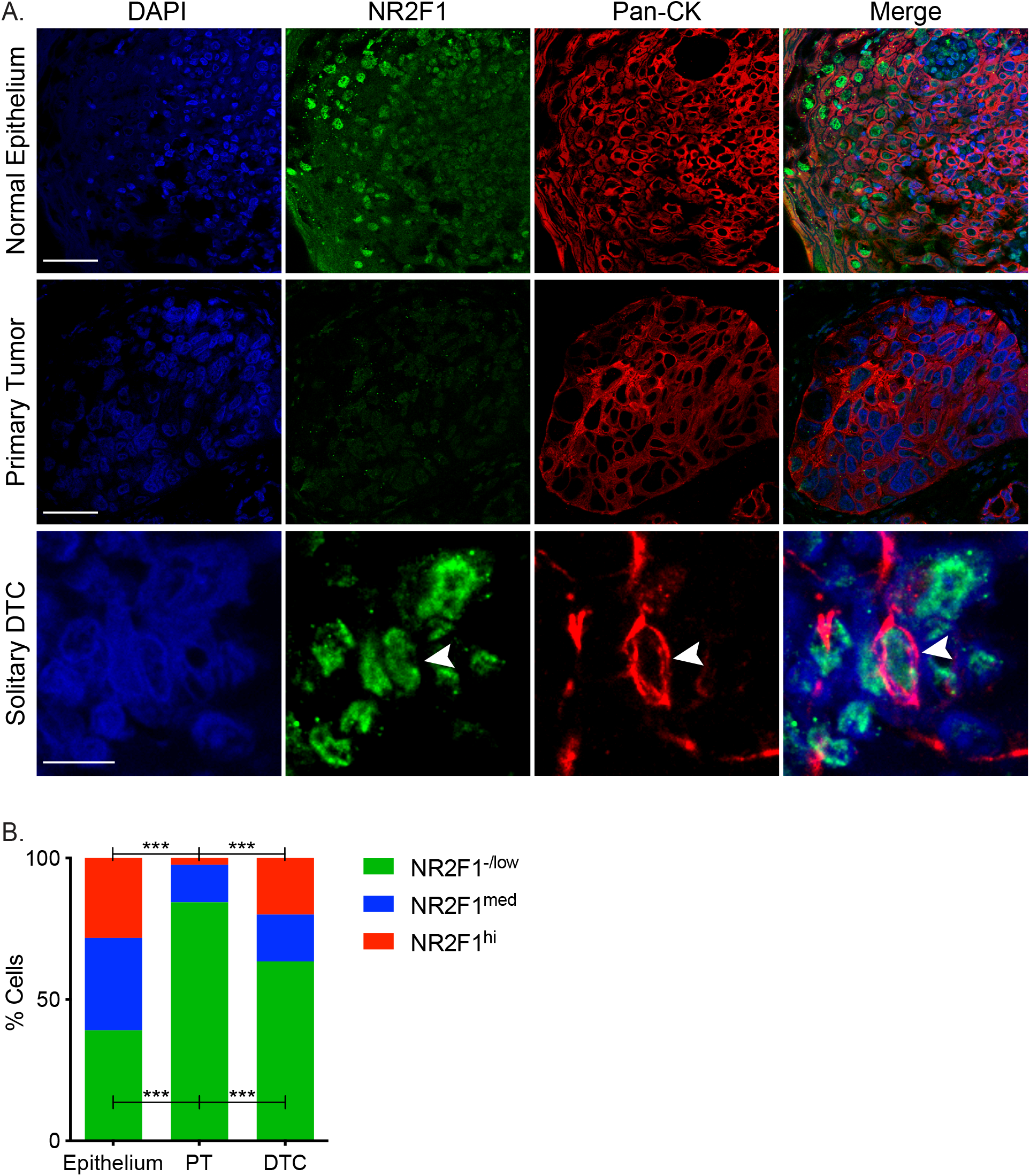
Solitary DTCs in HNSCC patients express high level of NR2F1. **A.** HNSCC Patient samples of normal epithelium, primary tumors, or lymph nodes with solitary DTCs were immunostained for NR2F1 and pan-cytokeratin (pan-CK) and the nuclei counterstained with DAPI. Scale bar for normal epithelium and primary tumor: 50 μm; scale bar for solitary DTC: 10 μm. **B.** Graph shows percentage of NR2F1^hi^, NR2F1^med^, or NR2F1^−/low^ cells in normal epithelium (n= 1814 cells from 2 patients), PT (n=3841 cells from 2 patients), or solitary DTCs (n=271 cells from 3 patients). *** p<0.001 by Fisher’s exact test adjusted with False Discovery Rate correction using Benjamini-Hochberg.

After establishing that upregulation of NR2F1 could mark dormant DTCs in patients, we wanted to verify that single DTCs in C26-treated mice are non-proliferative dormant cells. To this end, we immunostained lungs, which were collected from the previously described experiment, for Ki-67 and determined the percentage of proliferative Ki-67^+^/Vimentin^+^ tumor cells in lungs. While 45% of Vimentin^+^ tumor cells in DMSO control mice are Ki-67^+^, this number is significantly reduced to 7% of Ki-67^+^ in C26-treated mice (**Fig.6A and 6B**). Importantly, more than 99% tumor cells in the lungs of both DMSO and C26-treated mice were negative for cleaved caspase-3 (**Fig.S5A and S5B**), indicating that solitary DTCs in C26-treated lungs are not apoptotic. Collectively, these data show that single DTCs in lungs of C26-treated mice are indeed in a quiescent non-proliferative state. To determine if these single DTCs are in a dormant state, we immunostained for NR2F1, which not only is the target of C26, but also serves as a dormancy marker and is more informative on time to metastasis than proliferation markers alone as we showed in breast cancer DTCs from patients [7]. C26 treatment significantly almost tripled the frequency of NR2F1^high^ DTCs in lung as evident by an increase in the percentage of NR2F1^high^ cells (**Fig.6A and 6C**) and MFI per nucleus, which also doubles compered to control DTCs (**Fig.6A and 6D**). Interestingly, NR2F1 expression in DTCs of C26-treated mice was significantly higher compared to solitary DTCs in DMSO treated mice (**Fig.S5C**). These data indicate that C26 treatment maintains high NR2F1 expression in DTCs, which keeps them in a dormant state and prevents their growth into overt metastases. We also performed a similar analysis for the CDK inhibitor p27, which is a target of NR2F1 and also serves as a complementary dormancy marker. The percentage of cells with nuclear accumulation of p27 is significantly increased in lung DTCs of C26-treated mice compared to DMSO control (**Fig.6E and 6F**). This is also accompanied by an increase in nuclear p27 MFI (**Fig.6E and 6G**). Altogether, these results confirm that C26 treatment prevents outgrowth of lung metastases by activating NR2F1, which upregulates its own expression as well as downstream targets, including p27, and keeps DTCs in a dormant state.

**Figure 6.**
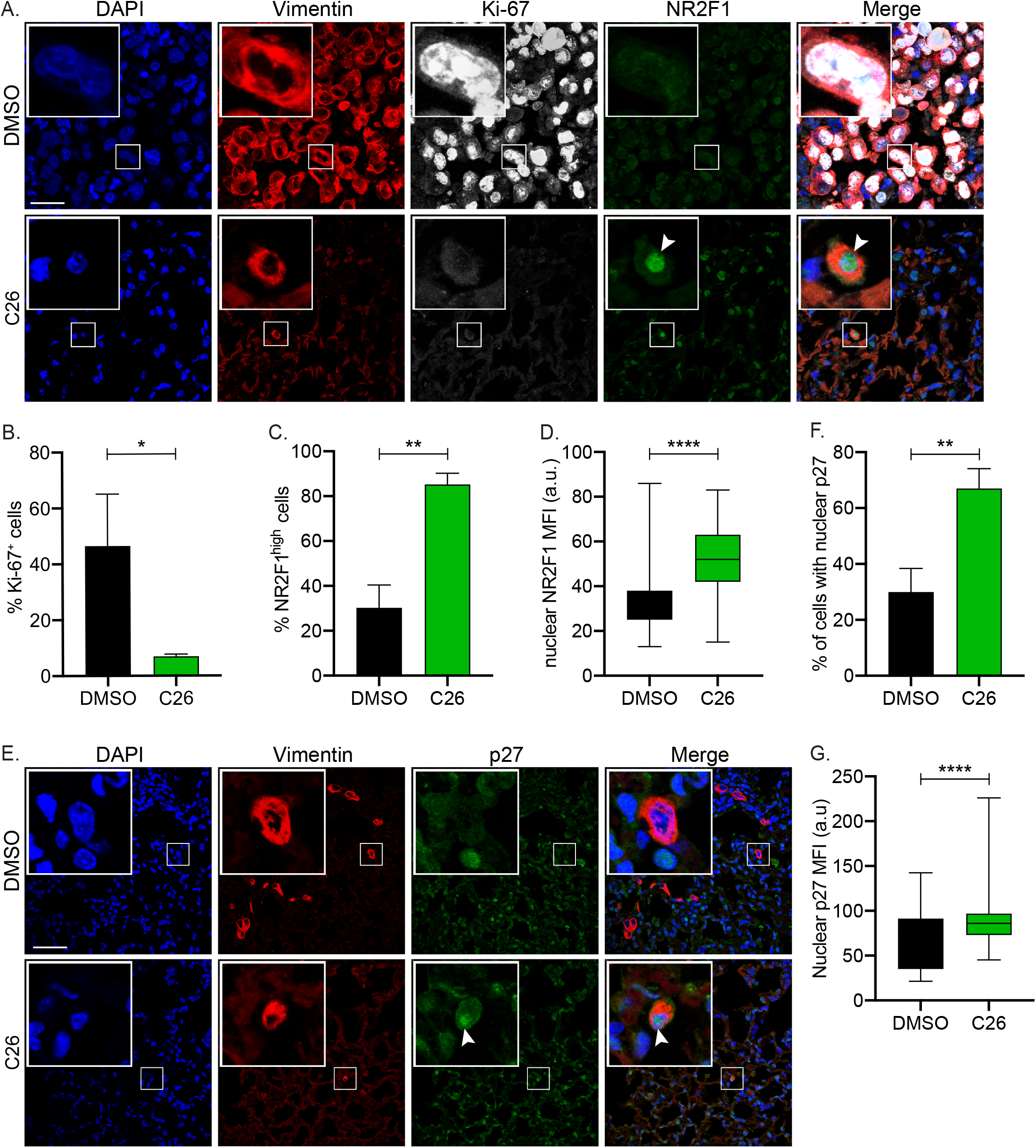
C26 suppresses metastatic growth in lungs by inducing dormancy. **A.** Lungs from DMSO or C26 treated mice described in Fig.4 were immunostained for vimentin, Ki-67, and NR2F1. Scale bar: 50 μm. **B.** Graph shows percentage of Ki-67^+^/vimentin^+^ tumor cells in lungs. **C.** Graph shows percentage of NR2F1^+^/vimentin^+^ tumor cells in lungs. Data in B and C represent mean +/− SEM. **D.** Graph shows box (25 to 75 percentile) and whiskers (minimum to maximum values) of nuclear NR2F1 mean fluorescence intensity (MFI) per cell in vimentin^+^ tumor cells in lungs from DMSO or C26-treated mice. Data in B, C, and D are from 5 mice per group (250 total cells for DMSO and 150 total cells for C26). * p<0.05; ** p<0.01; **** p<0.0001 by *t test*. **E.** Lungs from DMSO or C26 treated mice described in Fig.4 were immunostained for vimentin and p27. Scale bar: 50 μm. **F.** Graph shows percentage of vimentin^+^ tumor cells in lungs with nuclear accumulation of p27. Data represent mean +/− SEM. **G.** Graph shows box (25 to 75 percentile) and whiskers (minimum to maximum values) of nuclear p27 mean fluorescence intensity (MFI) per cell in vimentin^+^ tumor cells in lungs from DMSO or C26-treated mice. Data in F and G are from 5 mice per group (174 total cells for DMSO and 85 total cells for C26). ** p<0.01; **** p<0.0001 by *t test*.

## Discussion

We have previously discovered that NR2F1 is an important regulator of dormancy [3], a finding that was independently confirmed [3, 6, 11–13]. Thus, we sought to determine if this marker of dormancy could also be drugged to take advantage of its stable quiescence-inducing function. Our results using a structure-based *in silico* screening approach leveraged a new agonist that could accomplish our objective of turning “on” the NR2F1 pathway in malignant cancer cells. The inability of C26 to activate RXR□ and the NR21 knockout controls support the selectivity of C26, however it is possible that a crystal structure of NR2F1 LBD, which as of now has not been solved yet, may yield more potent and selective agonists. A surprising finding was that C26 induced NR2F1 expression in HNSCC T-HEp3 cells, which show silencing of NR2F1 through repressive histone modifications [3]. The induction of endogenous NR2F1 by C26 suggests that this silencing is not as tight as previously expected and that, although not tested directly, engaging NR2F1 with C26 may allow it to rapidly remodel the repressive chromatin at promoters, turn on its own expression, and drive downstream signals for dormancy onset. This is interesting, as activating this orphan nuclear receptor may be useful not only in treating NR2F1^+^ tumors, but also those that express low levels of NR2F1. NR2F1 agonists might be particularly useful in breast cancer where NR2F1 expression is highly enriched in ER^+^ tumors. Interestingly, these are the patients that commonly show late relapse [6]. Thus, agonists for NR2F1 may be used as a therapeutic approach to suppress reawakening of dormant cells kept in that state by anti-estrogen therapies or even in TNBC and other breast cancer subtypes.

We also wondered whether *in vivo* treatment with an NR2F1 agonist would be sufficient to prevent awakening of DTCs, and if the effect is long-lived. Our *in vivo* experiments in the CAM avian PDX system and in mice support that the NR2F1 function that C26 activates is long-lived and a strong growth suppressive signal. While pre-treatment of T-Hep3 in culture with C26 did not significantly inhibit cell proliferation *in vitro* (data not shown), it did suppress growth one week later *in vivo* on CAM, an effect that was stronger with continuous treatment *in vivo*. These data support that perhaps a treatment schedule that activates NR2F1 might not need continuous treatment as it may operate as a program that is self-sustained similar to what we previously reported for 5-azacytidine and retinoic acid treatment [3]. In the mouse PDX experiments, the neo-adjuvant + adjuvant strategy was chosen based on prior data showing that the very aggressive T-HEp3 PDX model spreads soon after subcutaneous implantation and that metastases can grow almost simultaneously with the primary tumor [20]. Nevertheless, our results strongly support that C26 was able to hold these DTCs, which are highly efficient at initiating metastasis [23, 24], in a dormant state. This was supported by the Ki67^lo^/p27^hi^/NR2F1^hi^ profile that these DTCs displayed. Furthermore, none of the animals treated with C26 displayed micro or macro-metastasis, strengthening the significance of activating NR2F1 to prevent metastatic relapse. Even when analyzing only NR2F1 levels specifically in solitary DTCs, which are more frequently spontaneously high for NR2F1 [3], we found that C26 treated animals displayed DTCs with higher NR2F1 expression than control animals. This corroborates that C26 might be preventing metastasis initiation by maintaining a dormant solitary cancer cell state. Surprisingly, C26 inhibited growth of some primary tumors. This further highlights the power of using NR2F1 agonists as a treatment strategy not only in an adjuvant setting (M0 patients in the clinic), but also to co-target proliferative and dormant DTCs co-existing in stage IV (M1) patients.

Mechanistically, there are some open questions. For example, one specific NR2F1 antibody that was used in the Matrigel and CAM assays reveals speckled nuclear localization of NR2F1, in large areas of the nucleus of the size of nucleoli. This signal was highly induced upon C26 treatment. Another NR2F1 specific antibody, which was used for paraffin embedded lung tissues, reveals a more homogeneous nuclear signal. These differences may indicate that there are different nuclear NR2F1 pools that might exhibit differential spatial localization and function, which requires further analysis. For example, it will be important to determine if C26 is inducing the same promoter and enhancer occupancy that NR2F1 shows during development [9] or in cells that have endogenously high NR2F1 levels. Similarly, we have not tested if C26 stabilizes the protein and whether this adds to the long-term effects we observed *in vitro* and *in vivo*. Finally, studies on reprogramming strategies, which were first designed in a cancer cell-centric way, revealed that they were also modulating the immune system to favor tumor control [25]. We have so far not explored the contribution of non-T cell-dependent responses to the effect of C26 on metastasis in nude mouse PDX models.

One of our objectives is to provide proof of principle data that dormancy modulation can be exploited to find alternative strategies to prevent metastatic disease. Our experience using a combination of 5-azacytidine and retinoic acid to activate global dormancy programs [3] was a first step. This knowledge led to the development of an ongoing clinical trial that has as a goal to treat prostate cancer patients with biochemical relapse (NCT03572387). We have now advanced a step further by rationally designing a strategy based on the selective activation of NR2F1. Our data have revealed that a selective NR2F1 agonist suppresses aggressive metastasis in a HNSCC PDX model, showing that it is possible to exploit dormancy mechanisms as therapeutic strategies to prevent metastasis. There is skepticism in the industry setting on advancing on dormancy-inducing strategies because there is a paralyzing misconception that trials need to last for many years, which would be costly. Our clinical trial in prostate cancer is a clear example that it is not accurate to assume that readouts of dormancy therapies would take long to be obtained. One could envision many scenarios where an agonist such as C26 could be tested within months. For example, in HNSCC, window of opportunity trials to test biological response for further patient selection for treatment in the adjuvant setting could be designed by randomizing treatment between biopsy-based diagnosis and surgery. Overall, our study reveals a mechanism-based and rationally designed strategy to exploit NR2F1 activated dormancy as a therapeutic option to prevent metastatic relapse.

## Materials and methods

### Cell lines

Tumorigenic (T-HEp3) HEp3 cells were derived from a lymph node metastasis from a HNSCC patient as described previously [26] and kept as patient-derived xenografts on chick chorioallantoic membrane (CAM). Dormant D-HEp3 cells were obtained by passing T-HEp3 cells for more than 40 generations in vitro [26]. When cultured *in vitro*, these cells were passaged in Dulbecco’s Modified Eagle Medium (DMEM) supplemented with 10% of fetal bovine serum (FBS) and 100 U penicillin/0.1 mg/ml streptomycin. SQ20B, FaDu, and HEK293T cells were obtained from ATCC and grown in the same medium as T-HEp3 and D-HEp3 cells. Cell transfection was performed using Lipofectamine 3000 (Invitrogen) following manufacturer’s instructions.

### Reagents and antibodies

Compound 26 (C26; molecular formula C28H30N4OS), was purchased from ChemBridge (#6596020). *All trans* retinoic acid (atRA) was purchased from Sigma-Aldrich (#R2625). Human NR2F1 plasmid was a kind gift from Dr. Gilles Salbert (Rennes University, France). Primary antibodies used for western blot and immunofluorescence are listed in supplementary Table S1.

### *In silico* NR2F1 LBD modeling and agonist screen

The three-dimensional structure of NR2F1 LBD domain was predicted by comparative protein structure modeling based on its alignment with the structure of RXRα LBD, whose crystal structure in complex with 9-cis retinoic acid has been solved (PDB code 1fm6; [16]). This analysis was performed using the software MODELLER v9.10 [27]. The resulting NR2F1 LBD was used to screen for potential agonists from a library of 110,000 drug-like small molecules (ChemBridge, USA) using computational docking. Docking scores were generated using two different softwares (Autodock 4 (release 4.2.3) [28] and eHITS [29]). In both cases, the docking grid was centered on the position of the 9-cis retinoic acid in the RXRα structure. The target and the ligands were prepared using AutoDock Tools.

### Luciferase reporter assay

HEK293T or D-HEp3 cells were transfected with cDNA for reporter vectors where firefly luciferase expression is driven by retinoic acid response element (RARE) [30] or NR2F1 cis element (Signosis #LR-2029). Cells were co-transfected with *Renilla* luciferase as an internal control. H3K293T cells were also transfected with NR2F1 cDNA. Cells were treated with DMSO or C26 for 36 hours after which they were lysed. Both *Renilla* and firefly luciferase activities were measured using dual-luciferase reporter assay kit (Promega #E1910) following manufacturer’s instructions. Data were reported as firefly/*Renilla* luciferase activity.

### RXRα activation assay

RXRα activation by C26 was assessed using LanthaScreen™ time-resolved fluorescence resonance energy transfer (TR-FRET) RXR alpha coactivator assay (ThermoFisher Scientific #PV4797). Briefly, a mixture of fluorescein-coregulator peptide and terbium anti-glutathione-S transferase (GST) antibody was added to GST-conjugated RXRα ligand binding domain (LBD) with varying concentrations of C26. DMSO and 9-cis-retinoic acid (9-cis-RA) were used as 0% activation control and 100% activation control, respectively. The terbium label on the anti-GST antibody was then excited at 340 nm and the energy transferred to the fluorescein label on the coactivator peptide was detected as emission at 520 nm. Emission ration (ER) was then calculated using the following formula: Fluorescein Emission (520 nm)/Terbium Emission (495 nm). Percent activation was then calculated using the following formula: (ER _Sample_ - ER _0% activation control_ / ER _100% activation control_ - ER _0% activation control_) x 100.

### Generation of knockout cell lines

NR2F1-knockout cell lines were generated using CRISPR/Cas9-targeted genome editing. Briefly, two separate NR2F1 guide RNAs (guide 2: 5’-GATCCGCAGGACGACGTGGC-3’ and guide 4: 5’-GGCTGCCGTAGCGCGACGTG-3’) were cloned into pLentiCRISPRv2 (Addgene #52961). A non-targeting (NT) guide RNA (5’-GTATTACTGATATTGGTGGG-3’) was used as a control. Lentiviral vectors were produced in HEK293T cells and viral supernatants were used to infect T-HEp3 or D-HEp3 cells. Cells were then selected using puromycin (Sigma-Aldrich #P8833). NR2F1 knockout was confirmed by western blot.

### 3D Matrigel assay

1000 cells (T-HEp3, FaDu, or SQ20B) were plated in 50 µl of growth factor-reduced Matrigel (Corning #356231) in 8-well chamber slides (Falcon). Cells were grown in media with reduced FBS content (2-5%). Cultures were treated every 24 hours starting at day 0 with DMSO or 0.5 µM C26. Single cells and clusters were counted after 4 days using a light microscope, and the cultures were then fixed for further analysis.

### Chick chorioallantoic membrane (CAM) assay

T-HEp3 tumor growth on CAM was previously described [31]. Briefly, 150×10^3^ T-HEp3 cells were inoculated on CAM and allowed to grow. Tumors were treated every 24-48 hours with 50 µl DMSO or 0.5 µM C26 starting at day 0. After 7 days, tumors were harvested and digested with collagenase-1A (Sigma #C9891) for 30 min at 37°C. T-HEp3 tumor cells, recognized by very large size compared to chicken cells, were counted using a hemacytometer.

### Mouse xenograft studies

750×10^3^ cells were injected subcutaneously in 8-week-old female BALB/c nu/nu mice (Jackson Laboratories) in the interscapular region. Mice were inspected every 48 h and arising tumors were measured with calipers in two perpendicular diameters. When tumors reached ~300 mm3, mice were treated intra-peritoneally with DMSO or C26 (0.5 mg/kg/day) for 5 consecutive days. Following 2 days of rest surgery was performed where mice were then injected with anesthetics ketamine 80–120 mg and xylazine 5 mg and a 1 cm incision was applied and the tumors were removed and weighed. Sutures were then performed using a wound clipper. After surgery, mice were subjected to 4 cycles of adjuvant therapy (5 days of treatment followed by 2 days of rest) with DMSO or C26 (0.5 mg/kg/day). At the end of the last cycle, mice were sacrificed and the lungs were collected and fixed in 10% formalin. All experimental procedures in mice were approved by the Institutional Animal Care and Use Committee (IACUC) of Icahn School of Medicine at Mount Sinai.

### Immunofluorescence

Matrigel cultures were fixed in 2% paraformaldehyde for 20 minutes and permeabilized in 0.5% TritonX-100 for 10 minutes. Paraffin-embedded sections were incubated in xylene followed by graded ethanol rehydration. Antigen retrieval for mouse lung tissues was performed in 10 mM citrate (pH 6) for 40 minutes using a steamer. Antigen retrieval for human tissue samples was performed in a steamer for 60 minutes in 1mM EDTA buffer. All paraffin-embedded tissues were permeabilized in 0.1% TritonX-100 for 10 minutes. Cells dissociated from CAM tumors were fixed in 4% paraformaldehyde, and cytospins were prepared by centrifugation onto glass slides. Cytospins were then permeabilized in 0.5% TritonX-100 for 10 minutes. Sections, cytospins, Matrigel cultures were blocked with 3% bovine serum albumin (BSA, Fisher Bioreagents) and 5% normal goat serum (NGS, Gibco #PCN5000) in PBS for 1 hour at room temperature. Primary antibodies were incubated overnight at 4°C followed by washing and incubation with alexa-conjugated secondary antibodies (Invitrogen, 1:1000 dilution) at room temperature for 1-3 hour in the dark. Slides were mounted with ProLong Gold Antifade reagent with DAPI (Invitrogen #P36931). Images were obtained using Leica Software on a Leica SPE confocal microscope and analyzed using ImageJ software.

### Western blot

Nuclear fractions were extraction using NE-PER Nuclear and Cytoplasmic Extraction Kit (Thermo Scientific #78835) and boiled for 5 minutes in sample buffer. Samples were run on SDS– PAGE gels and transferred to PVDF membranes. Membranes were then blocked in 5% milk in TBS-T. Primary antibodies (Table S1) were incubated overnight at 4°C. HRP-conjugated secondary antibodies were used at room temperature for 1 hour. Western blot development was done using ECL (GE #RPN 2106) and GE ImageQuant LAS 4010.

### Quantitative PCR

RNA was extracted using RNeasy mini kit (Qiagen #74104). cDNA was produced using High-Capacity cDNA Reverse Transcription Kit (Applied Biosystems #4368814). Real time PCR was performed using PowerUP SYBR Green Master Mix (Applied Biosystems #A25741). Primers used are listed in supplementary Table S2.

### Human samples

Paraffin embedded sections from HNSCC primary tumors were obtained from the Cancer Biorepository at Icahn School of Medicine at Mount Sinai, New York, NY. Paraffin embedded tissue sections from lymph nodes biopsied from clinically diagnosed metastasis-free HNSCC patients were obtained from Dr. Karl Christoph Sproll at the Heinrich-Heine-University Düsseldorf, Düsseldorf, Germany. These patients presented small primary tumors, had no previous malignancy in the head and neck region diagnosed before, and had not been subject to previous treatment. Samples were de-identified and obtained with Institutional Review Board approval. IF analysis was done from a total of 5 patients (2 patients with primary tumors and matched normal epithelium and 3 patients with lymph node DTCs).

### Statistical analysis

To ensure reproducibility, *In vitro* experiments were repeated at least three times unless otherwise indicated. For CAM tumor growth analysis, a minimum of 7 tumors were analyzed per group. For CAM qPCR and immunostaining experiments, a minimum of 4 tumors were analyzed. For mouse experiments, a minimum of 10 mice per group were used for tumor growth and number of lung DTC, while a minimum of 5 mice per group were used for immunostaining analysis. Statistical analysis was performed on Prism software using unpaired t test, 2-way ANOVA with Holm Sidak’s multiple comparison test, or Fisher’s exact test with false discovery rate correction using Benjamini-Hochberg where applicable. A p-value <0.05 was considered significant.

## Acknowledgments

We thank the Aguirre-Ghiso and Sosa labs for helpful discussions. We also thank Ms. Anna Freund MD and Ms. Marianne Hölbling for their technical assistance in preparing the FFPE sections.This work was supported by grants from the National Institute of Health /National Cancer Institute (CA109182, CA216248, CA218024, CA196521 to J.A.A-G.), the Samuel Waxman Cancer Research Foundation Tumor Dormancy Program (to J.A.A-G.), BioAccelerate NYC – NYC Partnership Fund (to J.A.A-G. and M.S.S), and HiberCell, LLC (to J.A.A-G.). B.D.K. was funded by National Cancer Institute T32 grant (CA078207). A.R.N. was funded by Portuguese Foundation for Science and Technology (SFRH/BD/100380/2014). K.C.S., together with Nikolas H. Stoecklein, was funded by the Deutsche Krebshilfe (#109600). Computational work reported in this paper was supported by the Office of Research Infrastructure of the National Institutes of Health under award numbers S10OD018522 and S10OD026880. The content is solely the responsibility of the authors and does not necessarily represent the official views of the National Institutes of Health.

## Author Contributions

BDK designed, planned and conducted experiments, analysed data and wrote the manuscript; RS and MM conducted the computational screen for agonists; TR, SM, and ARM conducted the *in vitro* agonist screen and phenotype validation experiments; BM and KCS, contributed tissue specimens and HNSCC expertise; EFF, JFC, ARN and NK planned and conducted the in vivo mouse experiments and processing of tissues; CRT conducted the experiments with patients samples; MSS and JAAG jointly conceived the project, designed experiments, analysed data and wrote the manuscript.

## Supplementary Figure Legends

**Figure S1.**
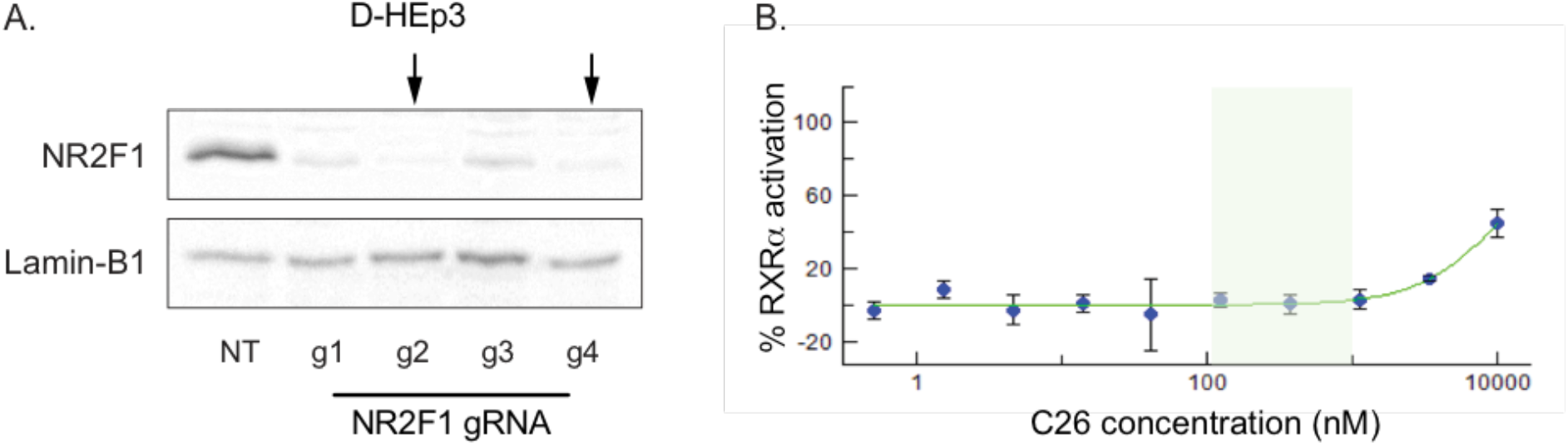
Supplementary data for Figure 1. **A.** Western blot shows NR2F1 protein expression level in nuclear extracts of D-HEp3 cell lines with non-targeting RNA (NT) or four different NR2F1 guide RNAs. Arrows indicate the two cell lines that were selected for use in experiments (g2 and g4). Lamin-B1 is used as a loading control. **B.** Graph shows percent activation of RXRα using different C26 concentrations. Shaded area represents the range of C26 concentration that was used in our experiments.

**Figure S2.**
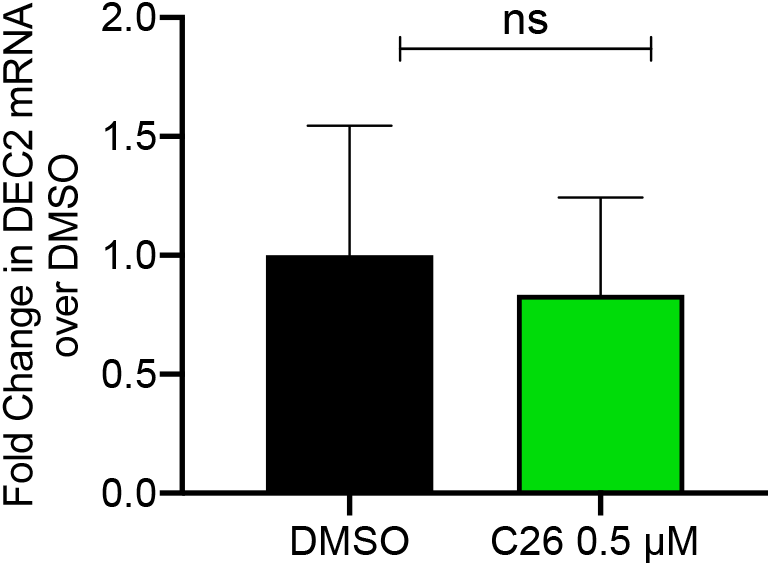
Supplementary data for Figure 2. DEC2 mRNA levels were measured using qPCR in DMSO or C26 treated CAM tumors. Graph shows fold difference in mRNA levels obtained by qPCR and normalized to DMSO. Data represent mean +/− SEM from 4 tumors per group. ns: not significant by *t test*.

**Figure S3.**
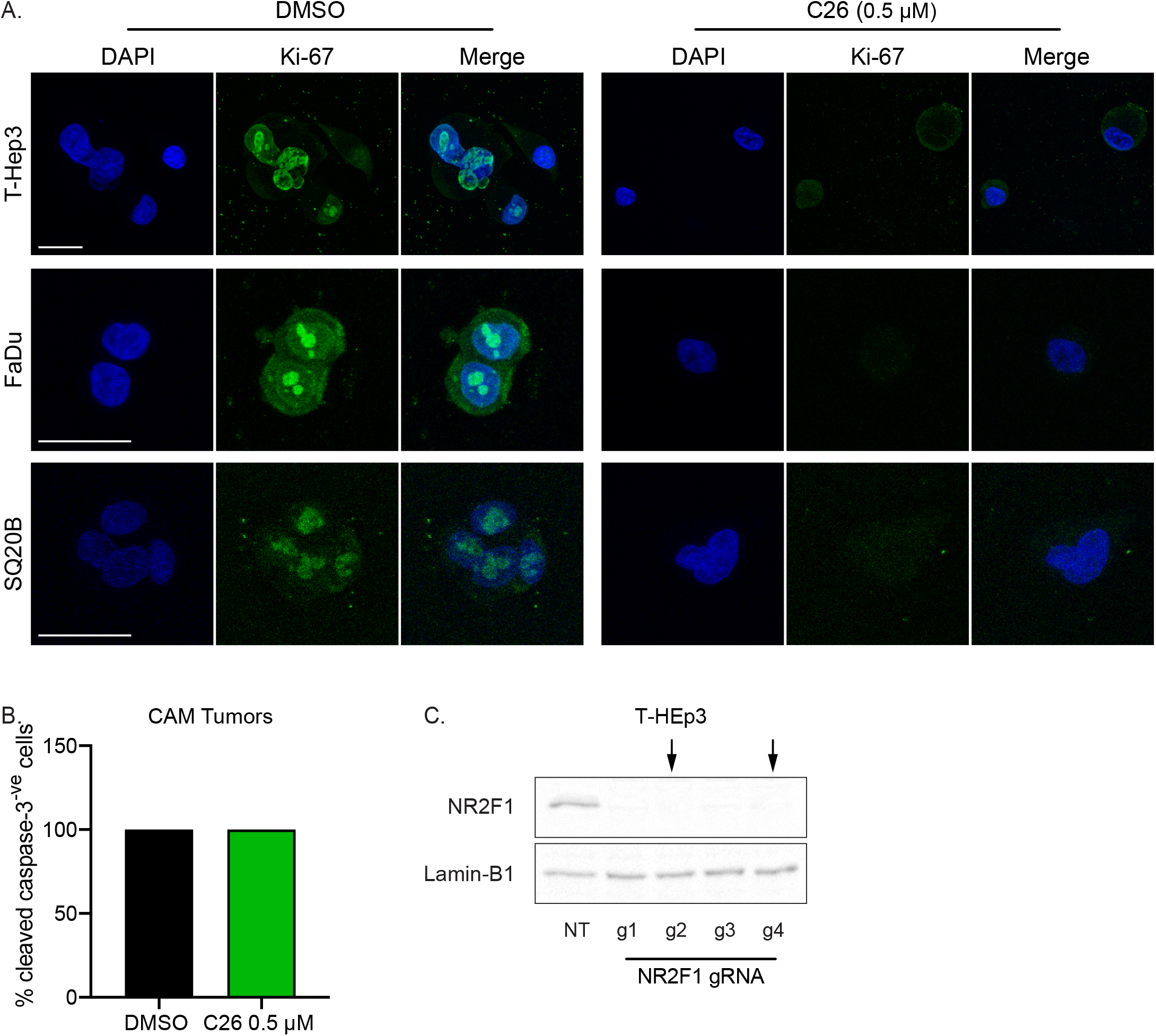
Supplementary data for Figure 3. **A.** Representative images of T-HEp3, FaDu, and SQ20B cells plated in Matrigel and treated with DMSO or C26 (0.5 μM) for 4 days then fixed and immunostained for Ki-67. Scale bar: 25 μm. **B.** DMSO or C26 (0.5 μM) treated CAM tumors were dissociated and cell cytospins were generated. Cytospins were immunostained for cleaved caspase-3 and nuclei counterstained with DAPI. Graph shows % cleaved caspase-3^+^ cells from 4 tumors per group. **C.** Western blot shows NR2F1 protein expression level in nuclear extracts of T-HEp3 cell lines with non-targeting RNA (NT) or four different NR2F1 guide RNAs. Arrows indicate the two cell lines that were selected for use in experiments (g2 and g4). Lamin-B1 is used as a loading control.

**Figure S4.**
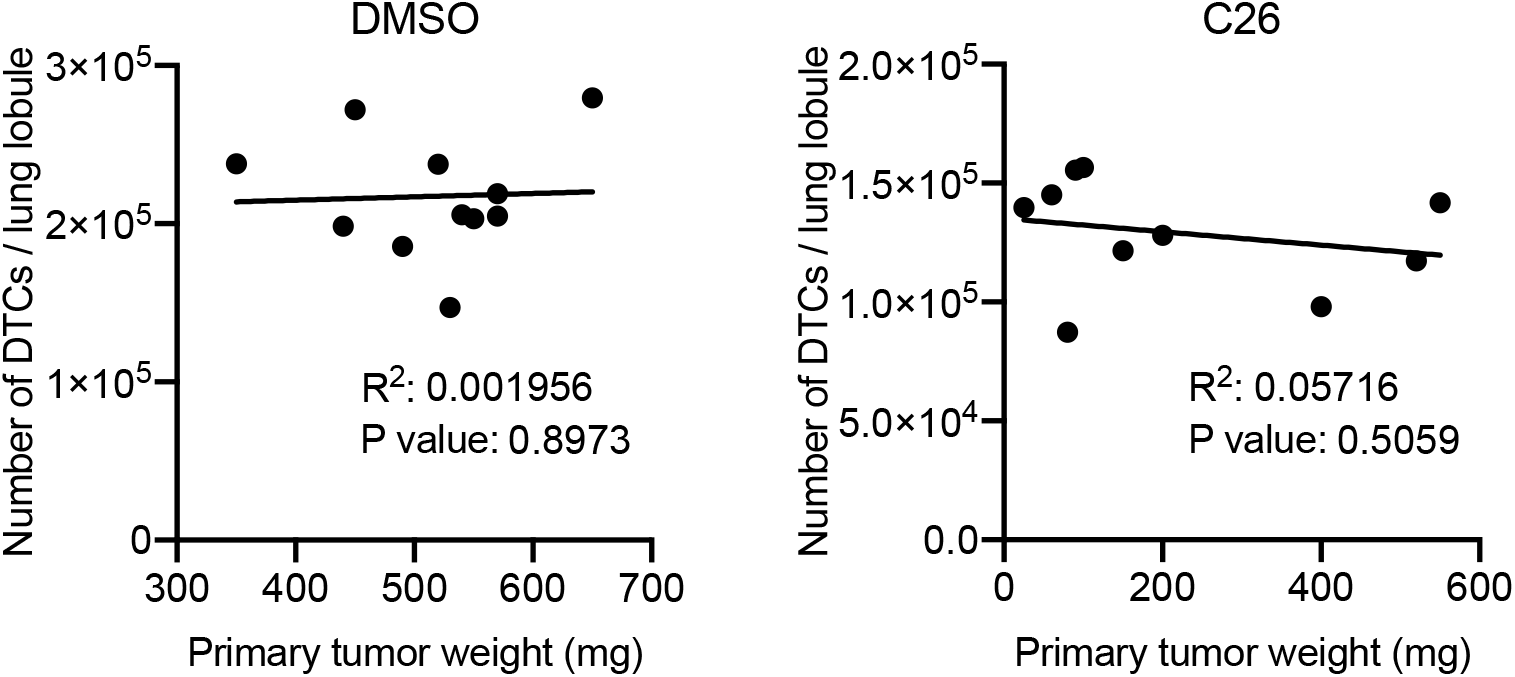
Supplementary data for Figure 4. Graphs show correlation between primary tumor weight and number of disseminated tumor cells (DTC) per lung lobule of DMSO or C26 treated mice.

**Figure S5.**
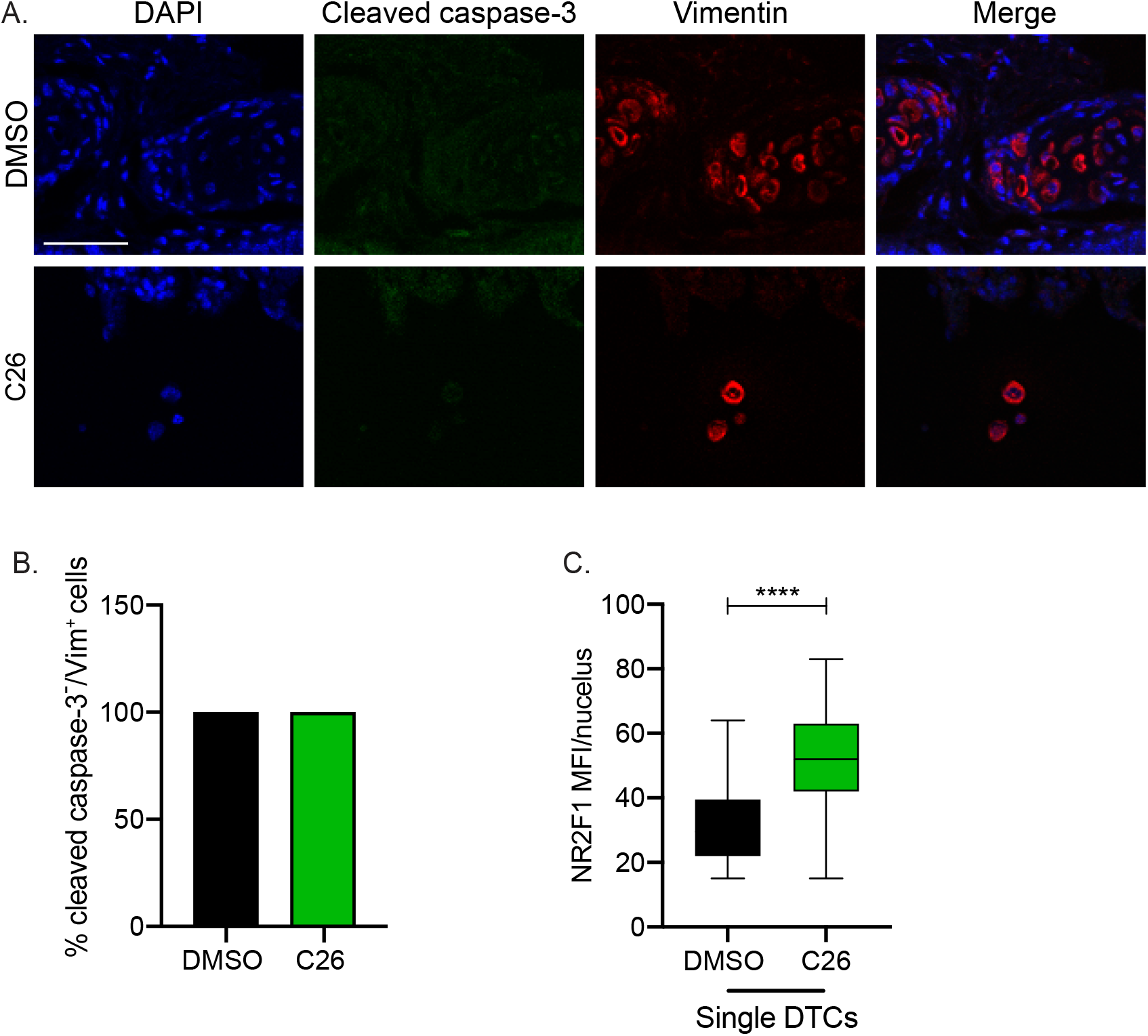
Supplementary data for Figure 6. **A.** Lungs from DMSO or C26 treated mice described in Fig.4 were immunostained for vimentin and cleaved caspase-3. Scale bar: 50 μm. **B.** Graph shows percentage of cleaved caspase-3^−^/vimentin^+^ tumor cells in lungs. **C.** Graph shows box (25 to 75 percentile) and whiskers (minimum to maximum values) of nuclear NR2F1 mean fluorescence intensity (MFI) in single DTCs only in lungs from DMSO and C26-treated mice. **** p<0.0001 by t test.

## Supplementary Table Legends

**Table S1.** Supplementary Table S1. List of primary antibodies used for IF and WB.

| Antibody | Manufacturer | Catalog number | Application | Dilution |
| --- | --- | --- | --- | --- |
| NR2F1 | R&D Systems | PP-H8132-00 | IF (Matrigel) | 1:100 |
|  |  |  | WB | 1:1000 |
| NR2F1 | Abcam | ab181137 | IF (paraffin sections) | 1:25-1:100 |
| Cleaved caspase-3 | Cell Signaling Technology | 9661 | IF | 1:100 |
| Ki-67 | Invitrogen | 14-5698-82 | IF | 1:100 |
| p27 | Cell Signaling Technology | 3686 | IF | 1:100 |
| Vimentin | R&D Systems | MAB2105 | IF | 1:100 |
| Lamin B1 | Abcam | 16018 | WB | 1:1000 |

**Table S2.**
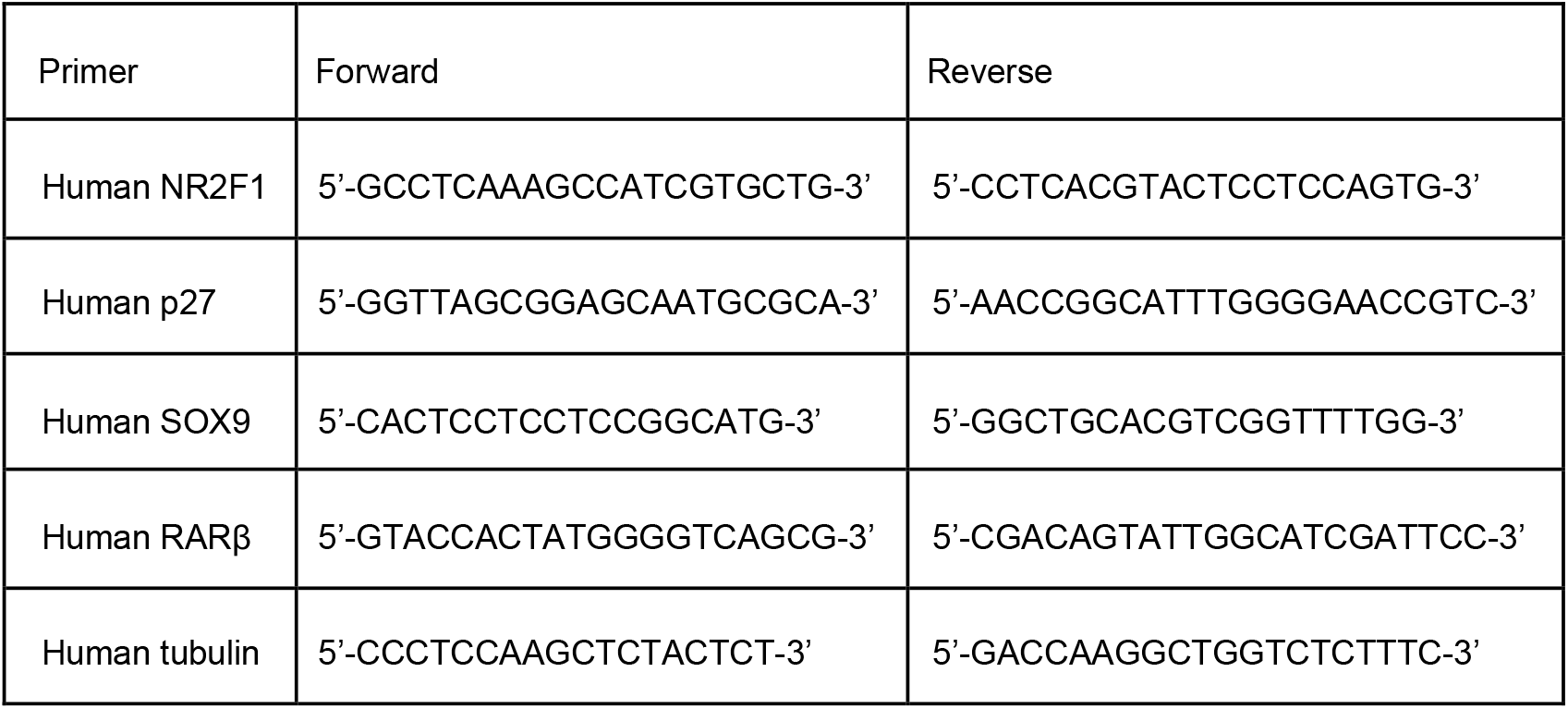
Supplementary Table S2. List of primers used for qPCR.

